# Differential levels of dermatan sulfate generate distinct Collagen I gel architectures

**DOI:** 10.1101/2020.05.05.078121

**Authors:** Jyothsna K M, Purba Sarkar, Keshav Kumar Jha, Varun Raghunathan, Ramray Bhat

**Affiliations:** Department of Electrical Communication Engineering, Indian Institute of Science, Bangalore, Karnataka 560012, India; Department of Molecular Reproduction, Development and Genetics, Indian Institute of Science, Bangalore, Karnataka 5600012, India; Leibniz Institute of Photonic Technology, Albert-Einstein-Straße 9, 07745 Jena, Germany

**Keywords:** collagen I, dermatan sulfate, second harmonic generation

## Abstract

Collagen I is the most abundant extracellular matrix (ECM) protein in vertebrate tissues. As an endogenously synthesized fibrillar biopolymer or as a synthetic hydrogel, it provides mechanical durability to tissue microenvironments and regulates cell function. Predominant regulators of its fibrillogenesis are dermatan sulfate proteoglycans (DSPGs), proteins conjugated with iduronic acid containing DS glycosaminoglycans (GAGs). Although DS is known to regulate tissue function through its modulation of Collagen I architecture, a precise quantifiable understanding of the latter remains elusive. We investigate this problem by visualizing the pattern of structural elements within fixed Collagen I gels polymerized in the presence of varying concentrations of DS (50-, 200- and 400- μg/mL) using second harmonic generation microscopy (SHG). Measuring four independent imaging parameters: fibril density, mean SHG signal (which estimates the ordering of the fibrils), surface occupancy (which estimates the space occupied by fibrils), and the fibril width allows us to construct an informative model of the effects of DS on Collagen I element architecture. Supported by confocal microscopy, our observations indicate that the effect on collagen fibril pattern of DS is contextual upon its concentrations. Lower levels of DS result in more numerous disorganized fibrils; higher levels restore organization, but at lower fibril densities. Such Collagen I gel pattern-tuning of DS is likely of relevance for understanding its functions in disease progression and biomaterial applications.

## INTRODUCTION

Collagen is the most abundant constituent of extracellular matrix (ECM), which is an essential component of the microenvironment of a functional tissue [1]. Apart from contributing to tissue function, it provides structural integrity and mechanical durability to physiologically functional tissues and organs [2, 3]. Within the family of collagens, the most prolifically expressed Type 1 Collagen or Collagen I polymerizes into fibrillar scaffolds in which cells of tissues are embedded [1, 4]. The fibril architecture of Collagen I can be modified by sulfated proteoglycans, which are proteins with long iterative chains of sulfated disaccharides known as glycosaminoglycans (GAGs). The expression and secretion of proteoglycans are tissue-specific; therefore proteoglycans contribute to making ECM distinct and specific to the tissues that secrete and remodel them [5, 6]. The largest class of proteoglycans comprises small leucine-rich proteoglycans (SLRPs) [7]. SLRPs are known to regulate collagen fibrillogenesis [8, 9]. They modulate collagen growth and organization of fibrils and fibers, extracellular matrix assembly, and by extension cell-matrix interactions, which are quintessential for cellular and tissue homeostasis [10]. GAGs have for long been used also in the studies involving the use of biomaterials and implants which often use hydrogels, including but not limited to Collagen I, where their ability to modulate hydrogel architecture and their biocompatibility are of immense benefit [11-13].

Proteoglycans containing GAGs known as dermatan sulfates (DS) are differentially expressed during developmental, physiological and pathological contexts [7]. DS consists of iterative units of *N*-acetyl-galactosamine (GalNAc) and iduronic acid (IdoA) [14, 15]. DS is the most abundant of GAGs that is associated with Collagen I and has been proposed to act by changing the geometry of the latter within stromal spaces [16]. Reese and coworkers polymerized Collagen I in the presence of DS and DSPGs. Using scanning electron microscopy as well as confocal reflectance microscopy, they found that DS increases the diameter of collagen fibrils [9]. On the other hand, Douglas and coworkers have observed a thinning of Collagen I and II fibrils upon addition of GAGs including DS [17]. The contradiction in the literature adds to the need of a precise quantitative demonstration of how and whether DS acts as a tuner of collagen gel architecture. This demonstration is crucial to the elucidation of mechanisms underlying the collagen related etiopathology of several diseases. Classic examples include congenital disorders such as the Ehler Danlos syndrome, spondyloepimetaphyseal dysplasia, San Filippo disease and several cancers where the DS or its conjugated protein (such as decorin, and biglycan) are upregulated [18-22]. Contemporary advances in microscopic modalities can provide fresh insights into physiological and pathological dynamics of fibrillar ECM topologies.

One such modality, second harmonic generation imaging is a well-established nonlinear optical contrast mechanism for imaging biological samples [23-25]. Demonstrated in biological samples first in 1986 [26], it is based on the upconversion of two lower energy photons to twice the incident frequency. Second harmonic signal is observed from objects that lack a center of symmetry, such as fibrillar extracellular matrix proteins and cellular cytoskeleton [27]. The associated chirality and non-centrosymmetry of collagen along with its highly ordered arrangement (tropocollagen assembling into fibrils, which in turn assemble into fibers) makes it an exceptionally strong source of SHG [28, 29]. Unlike fluorescence microscopy, the strong SHG signal from collagen allows label-free imaging of tissue samples without causing phototoxicity or photobleaching [30]. Moreover, it can be used to image structures in their native state in contrast with electron microscopic methods, which require sample dehydration or freezing. Such non-native state imaging can show significantly altered sample organization when compared to its native state. Since it is a nonlinear optical process involving two excitation photons, its axial and lateral resolution is higher (typically few hundred nanometers) when compared to diffraction limited resolution of single excitation photon [24].

Owing to these advantages, SHG microscopy has been used as a potential tool for analyzing changes in the collagen morphology in many diseased states [28]. SHG images combined with image analysis tools have been used to glean useful quantitative information such as fibril width, contrast, correlation, energy and homogeneity using image transforms such as curvelet transform, Fourier Transform matrix and grey-level co-occurrence matrix (GLCM), which were successful in elucidating structural changes accompanying ovarian cancer development [31, 32]. Above all, polarization resolved SHG metrices such as collagen peptide pitch angles, SHG signal anisotropy, SHG circular dichroism have also been used to classify ovarian cancer tissues at different stages [33].

In this manuscript, using SHG imaging we have studied the effect of DS on the gel architecture of Collagen I in detail. An integrative analysis of specific imaging parameters provides crucial insights into the significant alterations of element patterns by DS in a concentration-dependent and surprisingly, nonlinear fashion. We find that the lower levels of DS mold a Collagen I fibril pattern that is less ordered, with a greater number of shorter and thinner fibrils. On the other hand, the presence of higher levels of DS during polymerization lead to Collagen I fibrils that are ordered similar to control Collagen with relatively fewer long fibrils of equivalent width.

## METHODS

### I. Sample preparation

Acid extracted rat tail Type I Collagen (3 mg/mL; Thermo Fisher) was maintained at 4°C until polymerization. Dermatan sulfate sodium salt from porcine intestinal mucosa (Sigma-Aldrich) was reconstituted at 5 mg/mL. For collagen scaffolds, Type I collagen was neutralized in high salt conditions (10X DMEM) using 2N NaOH. Appropriate volume of DS GAGs were then added to neutralized collagen such that the final ratio of collagen: DS was 20:1, 5:1 and 2.5:1, referred to as DS (50 μg/mL), DS (200 μg/mL) and DS (400 μg/mL). Phosphate buffered saline was used to adjust the final volume. The mixtures were then plated onto chambered slides and incubated for 24 hours at 37°C in a carbon dioxide incubator for *in vitro* polymerization (See Figure 1A). At the end of 24 hours, the samples were fixed with 4% paraformaldehyde at 4°C for 30 minutes, following which they were washed with PBS thrice. The fixed sample was then taken for imaging using SHG and confocal microscopy.

**Figure 1.**
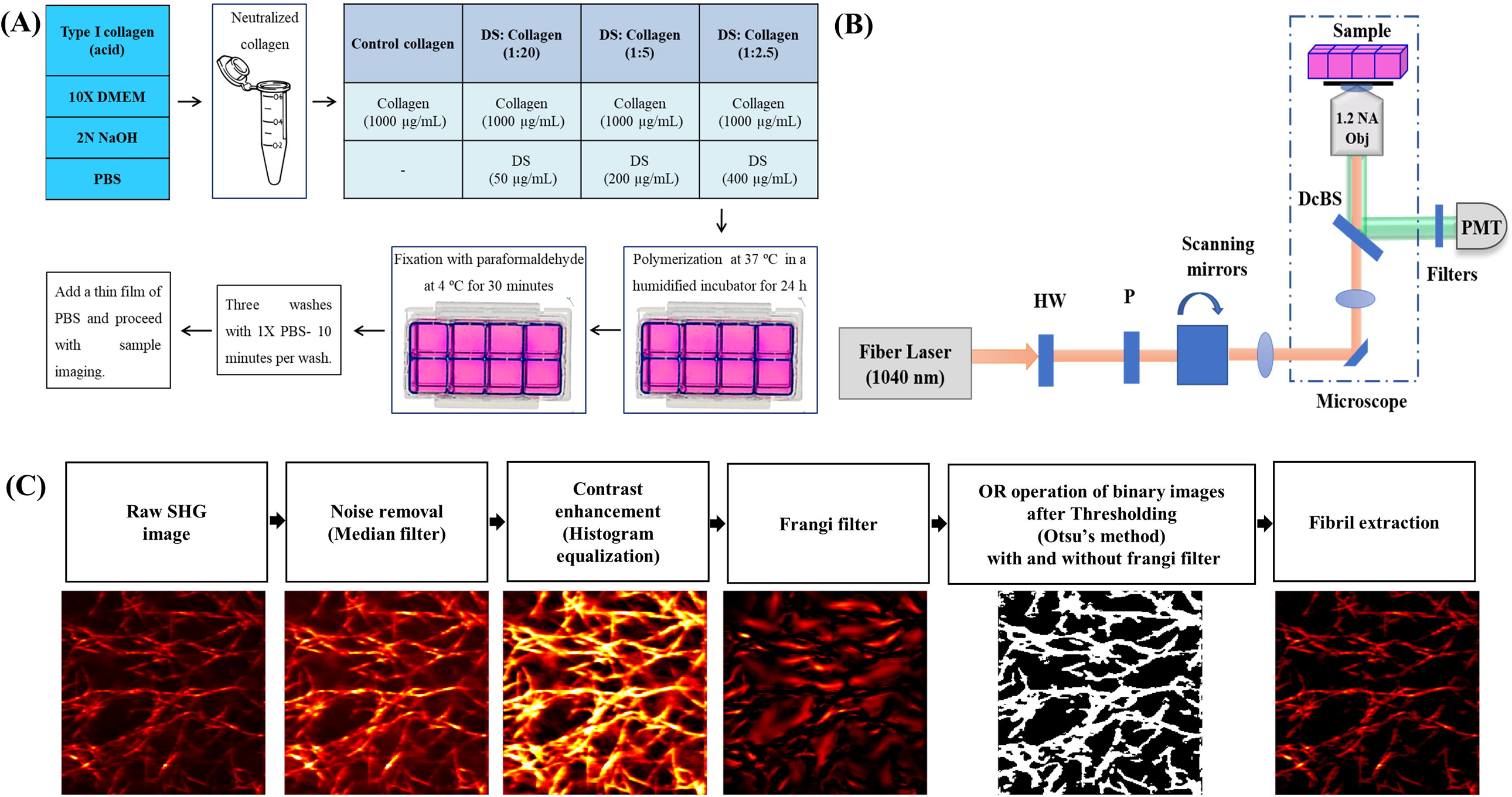
Description of sample preparation, imaging and analysis. (A) Flow chart depicting the *in vitro* polymerization of Collagen I in the absence and presence of Dermatan Sulfate followed by fixation and preparation for imaging studies (B) Schematic depiction of the second harmonic generation (SHG) microscopic set up used for sample imaging. HW: Half wave plate, PMT: Photomultiplier tube DcBS: Dichroic beam splitter, P: Polarizer and Obj: Objective (C) Flow chart showing the steps involved in fibril extraction using image processing approaches.

### II. Second harmonic generation and Confocal microscopy

A nonlinear microscopy set up shown in the Figure 1B is used for second harmonic generation imaging of the collagen polymerized in the presence and absence of DS. An optical source of fundamental excitation at 1040 nm by a femtosecond fiber laser (Fidelity HP) with 140 fsec pulse width and 80 MHz repetition rate is used as the incident light source at the fundamental frequency. Horizontally polarized fundamental excitation is focused onto the uniformly coated collagen gels in the chamber well mounted on an Olympus IX73 inverted microscope using a 60x/1.2 NA water immersion objective. The transverse and axial optical resolution at the SHG emission wavelength is estimated to be ∼ 467 nm and ∼858 nm, respectively. The second harmonic signal at 520 nm emitted from the sample is collected in epi-detection set up using the same objective. A dichroic mirror is used in the backward path to separate the fundamental and the second harmonic signal. The SHG signal is detected using a photomultiplier tube (Hamamatsu R3896) with a set of band-pass (520±20 nm) and short-pass (890 nm cut-off) filters in front to reject the fundamental excitation by ∼200 dB. SHG images of the collagen samples are obtained by scanning the laser beam using a pair of galvanometric mirrors (Thorlabs GVS002). More than 15 different sample locations or field of views (FOVs) of size 50 x 50 μm are imaged from each sample by moving the motorized sample stage (Thorlab MLS203-1). The pixel resolution used in the imaging is 200 nm. We restrict the image acquisition to a single, best focused image plane at every field-of-view. We acquired few images using a quarter wave-plate in the incident light path to create circularly polarized fundamental excitation and compared this with the linearly polarized excitation. The use of circularly polarized incident light did not result in any significant difference to the acquired SHG images.

For confocal microscopy, the collagen gels were polymerized and fixed on chambered glass slides as described in the sample preparation section. They were kept moist with PBS and imaged using a 63X oil immersion lens at the wavelength 633 nm. A total of 15 slices (FOVs) were imaged per stack using a 0.513 µm step size at a resolution of 1024 x 1024 pixels. All the confocal microscopy was performed using an inverted Zeiss LSM 880 with Airy scan.

### III. Image Analysis

The acquired images are processed in MATLAB and ImageJ to extract only the collagen fibrils from the images and used to calculate parameters such as mean SHG signal, surface occupancy, fibril width and the fibril density. The flowchart showing the various image processing steps used to identify the fibrillar structures in the Collagen images is shown in Figure 1C. Median filter is used to remove the noise and adaptive histogram equalization filter is used to enhance the contrast of the raw image. Frangi filter [34, 35] is applied onto the contrast enhanced image to sharpen the edges of the fibrils. Binarization is done on the contrast enhanced image with and without frangi filter using Otsu’s method. Otsu’s method is a non-parametric method that automates the optimal threshold calculation by using zeroth and first order cumulative moments of the gray-level histogram from the image [36]. The resulting binary images are added (OR operation) to retrieve all the fibril pixels (used to calculate surface occupancy), which is further multiplied with the raw image (used to calculate mean SHG) and the contrast enhanced image (used to calculate fibril width and density) to highlight selectively the individual collagens fibrils.

Mean SHG signal, which is a measure of the ordering of the structural elements of collagen across the imaged field-of-view is calculated from the above processed image by calculating the average SHG value after selecting only the image pixels corresponding to the collagen fibrils. Surface occupancy is defined as the ratio of the number of pixels occupied by the fibrils to the total number of pixels in the image. This is calculated from the binarized image by taking the ratio of number of white pixels to the total number of pixels in the binary image. Fibril density, which measures the number of fibrils within a given plane irrespective of their arrangement, and fibril width are estimated from the contrast enhanced images using Ridge detection, an ImageJ plugin [37]. After specifying parameters such as line width (approximate fibril diameter in pixels), highest and lowest possible grey scale value of the fibrils, this algorithm uses differential geometry-based approach for detecting the fibrils and their edges to extract the widths of the fibrils with sub-pixel resolution [38]. Trial and error approach is used to estimate the above-mentioned input parameters with best accuracy. The extracted fibril width in pixels is then multiplied with the appropriate conversion factor corresponding to the pixel resolution to obtain the fibril width in microns. The fibrils or structural elements with widths greater than 0.1 μm are counted to calculate the number of fibrils within a given plane, referred to here as fibril density.

### IV. Statistical analysis

This experiment was repeated four times and during each trial, we imaged more than 15 FOVs from each sample. All the data are presented with mean and standard deviation of the mean. Comparison between two groups is done using unpaired t test and repeated measures (RM) one-way ANOVA to compare the data of the four experiments. Any data set with p-value less than 0.05 is considered statistically significant. All the analyses were done using GraphPad Prism software. Table 1 summarizes the values of the estimated parameters for four sets of experiments with corresponding p-values.

**Table 1:** Comparison of quantitative parameters characterising the effect of DS on the fibrillar gel architectures of collagen I tabulating values for each parameter for four independent experiments and scores signifying statistical significance between their means.

| Parameters | N | Control<br>Collagen I | Collagen I-DS<br>(50 µg/mL) | Collagen I-DS<br>(200 µg/mL) | Collagen I-DS<br>(400 µg/mL) | P value<br>( RM One-way<br>Anova test) |
| --- | --- | --- | --- | --- | --- | --- |
| Mean SHG signal (V) | 1 | 0.522 | 0.267 | 0.430 | 0.458 | 0.0339 |
|  | 2 | 0.719 | 0.386 | 0.511 | 0.758 |  |
|  | 3 | 0.597 | 0.626 | 0.636 | 0.628 |  |
|  | 4 | 0.533 | 0.314 | 0.575 | 0.543 |  |
| Fibril density | 1 | 324 | 412 | 273 | 251 | 0.0162 |
|  | 2 | 443 | 490 | 489 | 328 |  |
|  | 3 | 481 | 505 | 441 | 435 |  |
|  | 4 | 331 | 350 | 320 | 301 |  |
| Fibril width (µm) | 1 | 1.445 | 1.280 | 1.390 | 1.433 | 0.0342 |
|  | 2 | 1.450 | 1.370 | 1.390 | 1.476 |  |
|  | 3 | 1.460 | 1.447 | 1.439 | 1.434 |  |
|  | 4 | 1.404 | 1.325 | 1.437 | 1.424 |  |
| Surface occupancy (%) | 1 | 54.55 | 41.57 | 46.83 | 45.86 | 0.0461 |
|  | 2 | 65.13 | 51.81 | 57.86 | 60.81 |  |
|  | 3 | 62.40 | 62.75 | 58.70 | 57.27 |  |
|  | 4 | 54.70 | 44.33 | 56.27 | 53.45 |  |

## RESULTS

### Incorporation of DS alters the ordering of Collagen I fibrils

Analysis of the effect of DS on the ordering/regularity of Collagen I fibril patterns was carried out by comparing the mean SHG signals. Mean pixel intensity and hence mean SHG signal is expected to be higher, when the fibrils are aligned (and even bundled) with respect to each other [39]. When examined, the fibrils of the control Collagen I appeared long and organized with parallel alignments in several fields, which is prognostic of bundling of fibrils into higher order structural elements like fibers (Supplementary Figure 1A). In comparison, fibrils of Collagen I-DS (50 μg/mL) showed a more disorganized web-like arrangement with much lower frequency of bundling. Collagen I-DS (200 μg/mL) and Collagen I-DS (400 μg/mL) fibrils were visually indistinguishable from control Collagen I and also shows a higher frequency of organized and often bundled appearance (Figure 2Ai-iv). It was therefore unsurprising that the difference in mean SHG, when compared across 4 independent biological experiments using repeated measures-ANOVA, was found to be significant (p-value = 0.0339), with the Tukey’s post hoc comparison showing significant difference between Collagen I-DS (50 μg/mL) on the one hand, and Collagen I-DS (400 μg/mL) and control on the other (Figure 2B).

**Figure 2.**
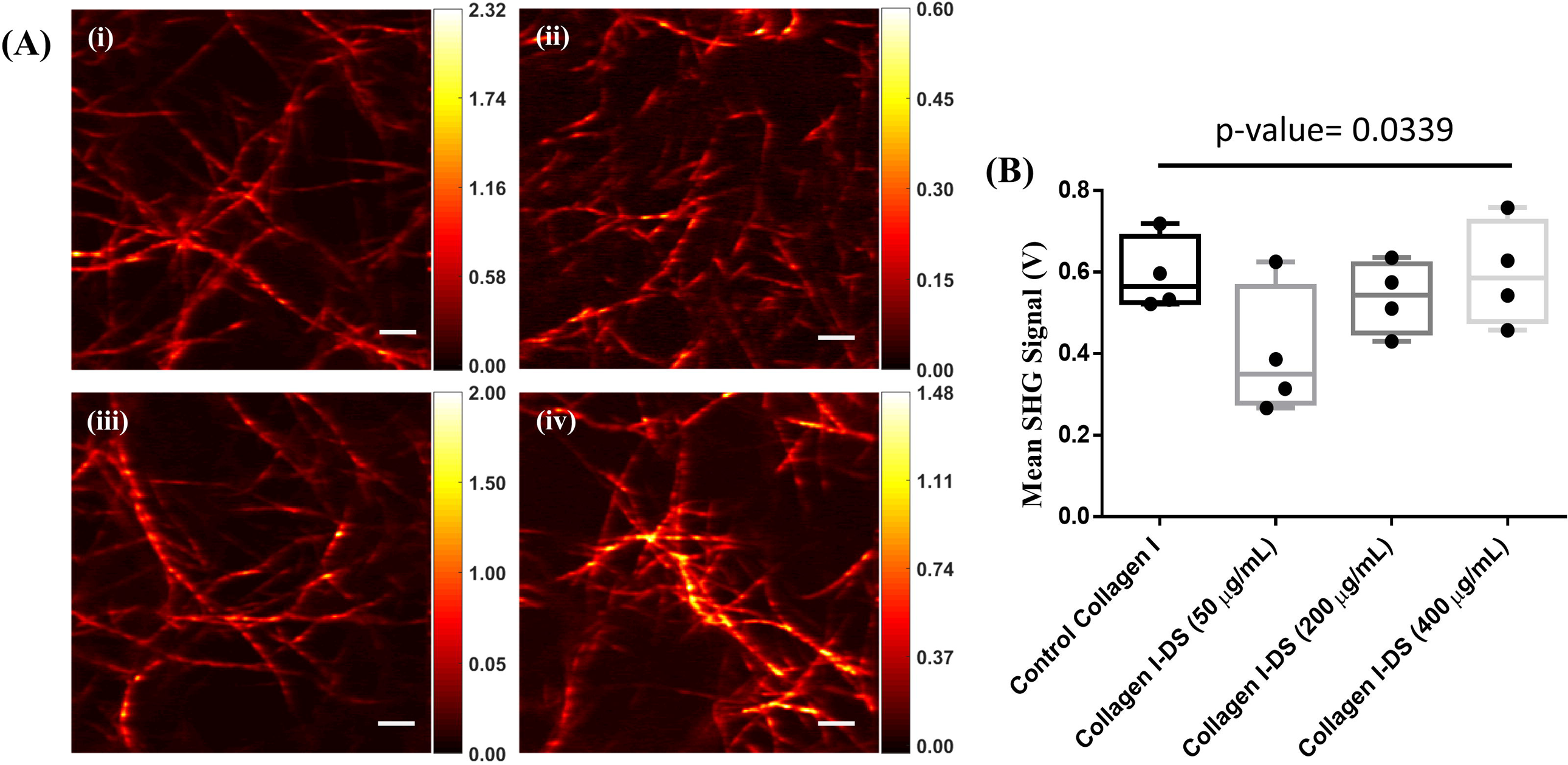
Effect of Dermatan Sulfate on mean SHG signal of collagen I fibril architecture. (A) SHG photomicrographs demonstrating the SHG signal from collagen I matrices polymerized in the absence of dermatan sulphate (i) and presence of dermatan sulphate at 50 µg/mL (ii), 200 µg/mL (iii) and 400 µg/mL (iv) respectively. Color bar labels show PMT signal in Volts and scale bar = 5 µm. (B) Bar graph showing the mean SHG signal under the conditions indicated in 2A. Statistical significance was computed using repeated measures ANOVA with Tukey’s post hoc analysis of difference in means. Error bars show the mean ± SD of the four experiments with more than 15 fields of view from each sample.

### Incorporation of DS in Collagen I alters the fibril density of the scaffolds

We next asked if the number of fibrils in a given space was altered by the presence of DS. To further probe this, we used the ridge detection algorithm to calculate the fibril density (number of fibrils) per FOV and its relationship with DS concentration during polymerization. A greater number of fibrils would therefore signify higher fibril density. To our surprise, the SHG images pertaining to Collagen I-DS (200 μg/mL) and Collagen I-DS (400 μg/mL) showed a lower density of fibrils when compared with Collagen I-DS (50 μg/mL), which on the other hand showed the highest fibril density among the four examined samples (Figure 3Ai-iv). Repeated measures ANOVA confirmed these observations (p value = 0.0162), although Tukey’s multiple comparisons test only showed a significant separation between Collagen I-DS (50 μg/mL) and Collagen I-DS (400 μg/mL) suggesting the maximum divergence in density between these two samples (Figure 3B). These findings may be reconciled with our observations on mean SHG in that lower DS levels seem to modulate fibril organization away from higher order elements resulting in an increase in their number. On the other hand, higher levels of DS mediate more bundling resulting in more higher-order elements (such as fibers) and lesser number of fibrils.

**Figure 3.**
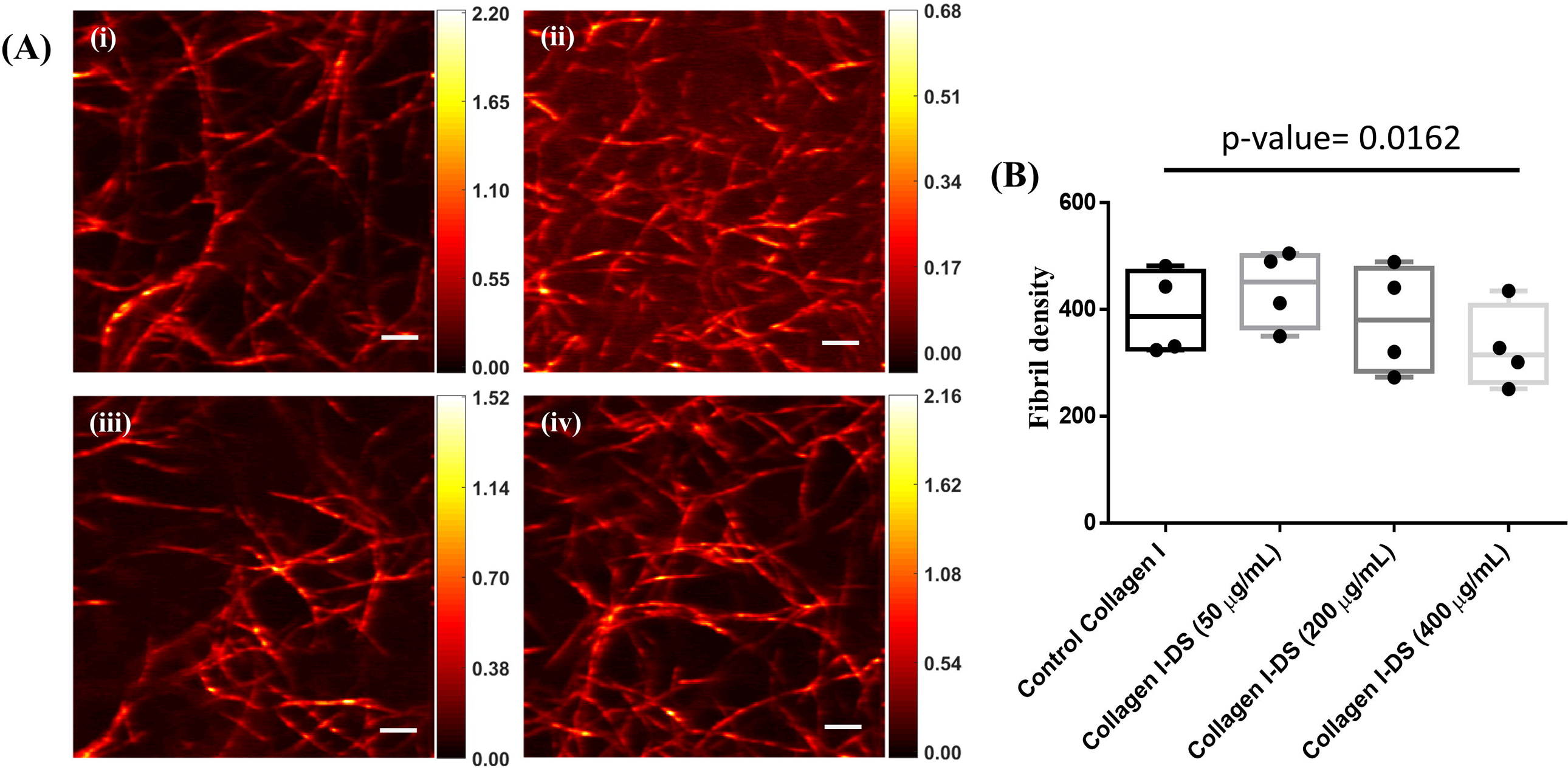
Effect of Dermatan Sulfate on density of collagen I fibril architecture. (A) SHG photomicrographs demonstrating the fibril density of collagen I matrices polymerized in the absence of dermatan sulphate (i) and presence of dermatan sulphate at 50 µg/mL (ii), 200 µg/mL (iii) and 400 µg/mL (iv) respectively. Color bar labels show PMT signal in Volts and scale bar = 5 µm. (B) Bar graph showing the fibril density under the conditions indicated in 3A. Statistical significance was computed using repeated measures ANOVA with Tukey’s post hoc analysis of difference in means. Error bars show the mean ± SD of the four experiments with more than 15 fields of view from each sample.

### Incorporation of DS in Collagen I alters Collagen fibril width

We next investigated whether DS also regulated the width of structural elements formed within polymerized Collagen I gels. SHG imaging revealed a mild decrease in fibril width when Collagen I was polymerized in the presence of DS (50 μg/mL) compared with control. Higher concentrations of DS (200 and 400 μg/mL) did not show any significant differences when compared with control (Figure 4Ai-iv). Repeated measures ANOVA with Tukey’s post hoc comparison served to confirm the significantly lower fibril width of Collagen I-DS (50 μg/mL) with respect to control Collagen I and Collagen I-DS (400 μg/mL) (p-value = 0.0342) (Figure 4B).

**Figure 4.**
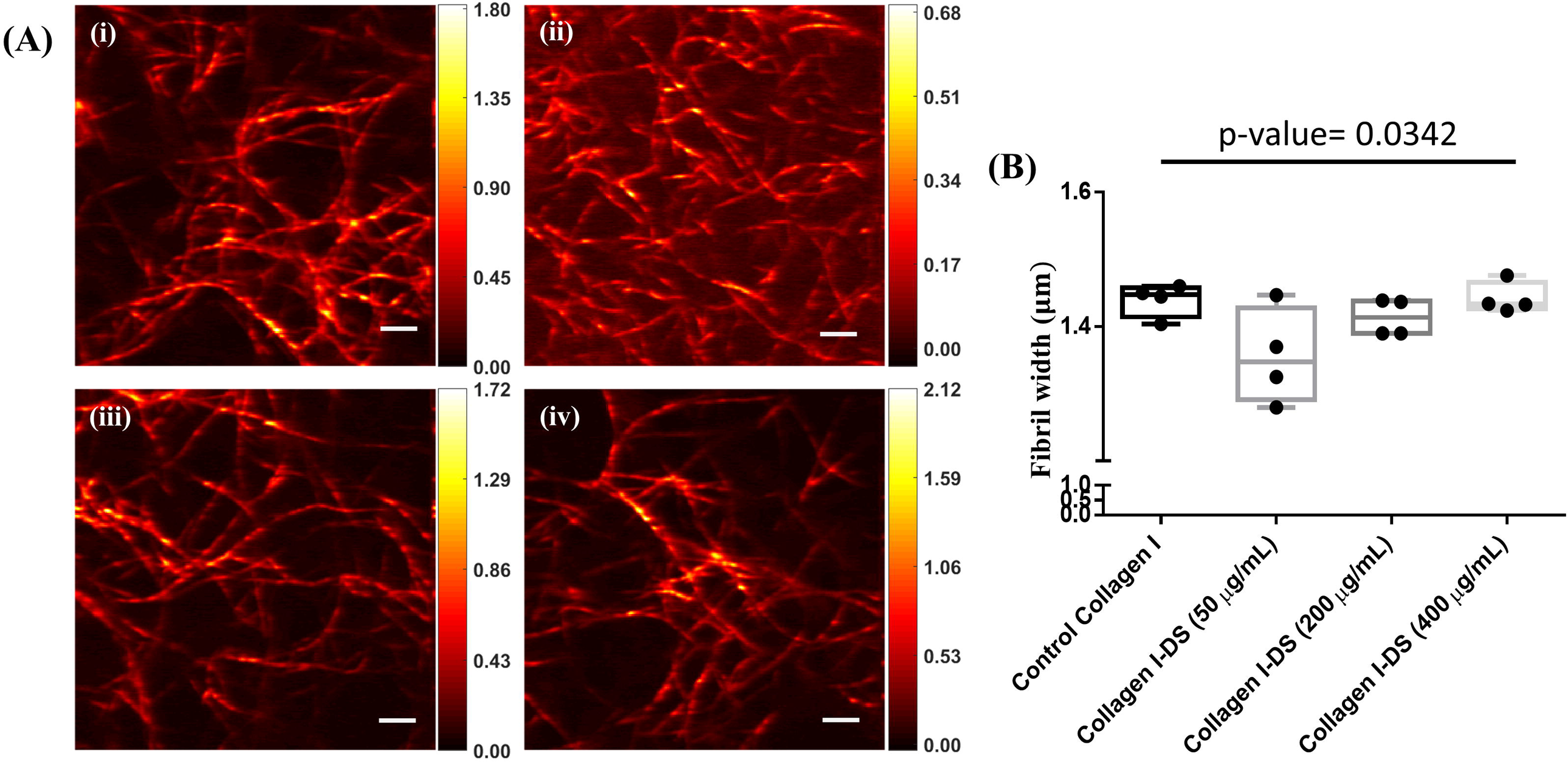
Effect of Dermatan Sulfate on width of collagen I fibrils within gel architectures. (A) SHG photomicrographs demonstrating the fibril width of collagen I matrices polymerized in the absence of dermatan sulphate (i) and presence of dermatan sulphate at 50 µg/mL (ii), 200 µg/mL (iii) and 400 µg/mL (iv) respectively. Color bar labels show PMT signal in Volts and scale bar = 5 µm. (B) Bar graph showing the fibril width under the conditions indicated in 4A. Statistical significance was computed using repeated measures ANOVA with Tukey’s post hoc analysis of difference in means. Error bars show the mean ± SD of the four experiments with more than 15 fields of view from each sample.

### Incorporation of DS in Collagen I alters the occupancy of the fibrils on the scaffold surfaces

The last parameter we estimated was surface occupancy, which describes the packing of the collagen fibrils in the given space or volume. When imaged, fibrils were found to occupy a greater proportion of the imaged plane in control Collagen I, Collagen I-DS (200 μg/mL) and Collagen I-DS (400 μg/mL) compared with Collagen I-DS (50 μg/mL) (Figure 5Ai-iv). The statistical significance of this observation was confirmed through quantitative comparison across independent biological experiments using repeated measures-ANOVA (p-value = 0.0461). (Figure 5B). Although the values for Collagen I-DS (200 μg/mL) and Collagen I-DS (400 μg/mL) on the one hand and Collagen I-DS (50 μg/mL) on the other, were distinctly separated, Tukey’s post hoc comparison indicated significance only between the latter and control Collagen I (Figure 5B).

**Figure 5.**
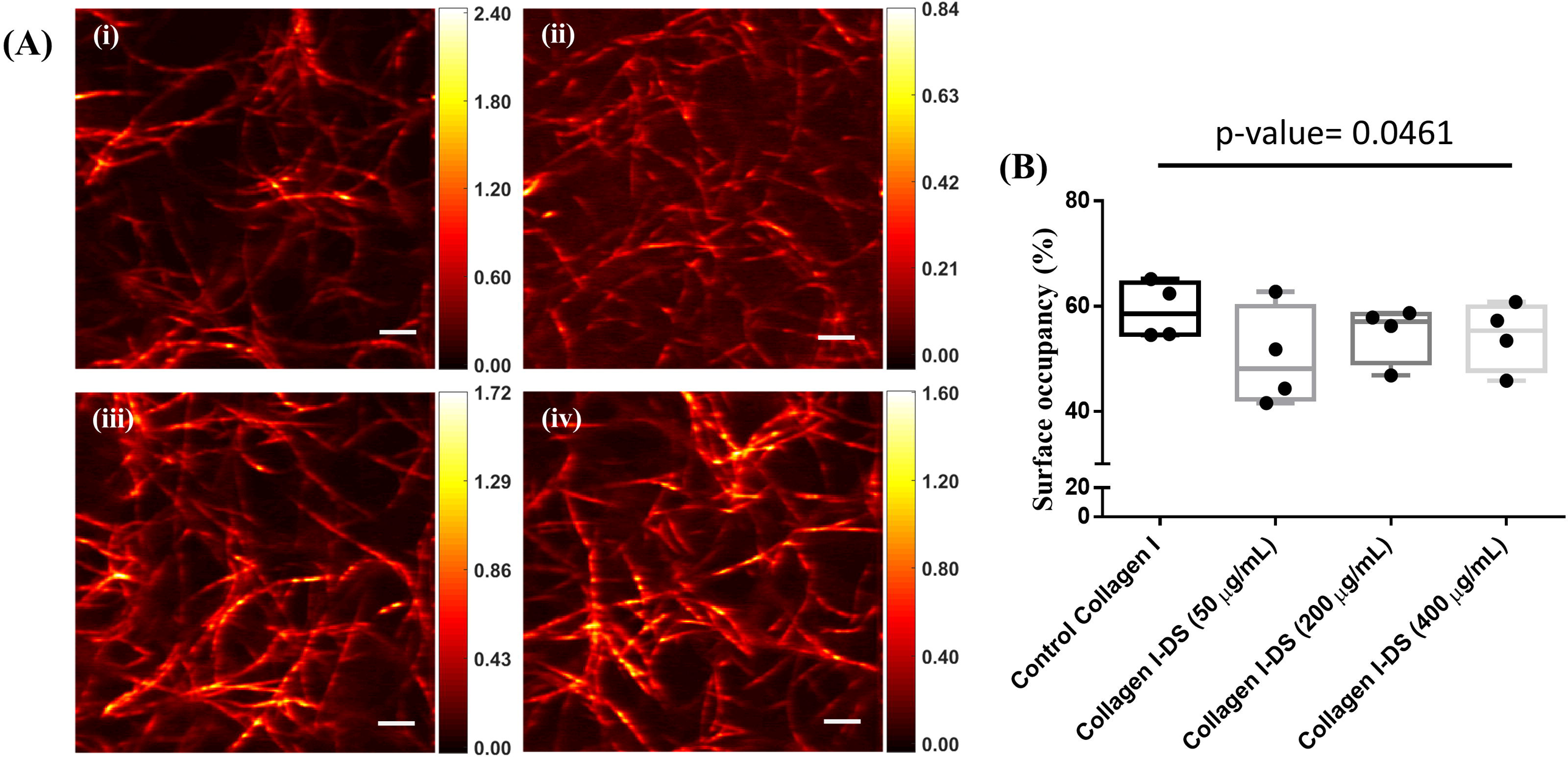
Effect of Dermatan Sulfate on surface occupancy of collagen I fibril architecture. (A) SHG photomicrographs demonstrating the surface occupancy (for definition, please see Materials and Methods) of collagen I matrices polymerized in the absence of dermatan sulphate (i) and presence of dermatan sulphate at 50 µg/mL (ii), 200 µg/mL (iii) and 400 µg/mL (iv) respectively. Color bar labels PMT signal in Volts and scale bar = 5 µm. (B) Bar graph showing the surface occupancy under the conditions indicated in 5A. Statistical significance was computed using repeated measures ANOVA with Tukey’s post hoc analysis of difference in means. Error bars show the mean ± SD of the four experiments with more than 15 fields of view from each sample.

### Change in pattern of Collagen I fibrils by DS imaged by SHG is consistent across fields of image acquisition

The calculation of the above-studied four imaging parameters were arrived at through acquiring and combining measurements across >15 fields of view of scaffolds for each independent biological experiment. We sought to confirm the veracity of the trends assimilated across the four experiments, by inferring them within each separate single biological experiment. The order of the parameter magnitudes (mean values) was similar for 3 of 4 independent experiments, with the addition of DS at 50 μg/mL correlating with lower fibril ordering, surface occupancy and fibril width and greater fibril density. Higher concentrations of DS (200- and 400 - μg/mL) on the other hand were able to restore fibril ordering, surface occupancy and fibril width to control levels and showed lower fibril density with respect to the latter. Supplementary Figure 2 shows the differences in means of all four parameters on one of the biological experimental combinations whereas Table 1 shows the parametric values obtained across all four experiments that were used to compute the cumulative statistics.

### Imaging of Collagen I using laser confocal microscopy supports the SHG microscopy-based results

To further confirm some of the results obtained from quantitative analysis done on SHG images using an independent linear optical method, we acquired confocal images of the reflectance signals emanated by control Collagen I, Collagen I-DS (50 μg/mL) and Collagen I-DS (400 μg/mL) gels. Figure 6 (A-C) shows the representative confocal images of control Collagen I, Collagen I-DS (50 μg/mL) and Collagen I-DS (400 μg/mL) respectively. For the same laser intensity, the reflectance signals from the Collagen I fibrils were stronger in the control Collagen I and Collagen I-DS (400 μg/mL) scaffolds than Collagen I-DS (50 μg/mL), which showed lower signals. Numerous long and parallel fibrils could also be appreciated in the control Collagen I sample (see also Supplementary Figure 1B). In Collagen I-DS (50 μg/mL) gels, a number of shorter fibrils could be appreciated. However, the Collagen I-DS (400 μg/mL) scaffold showed longer fibrils but lower in density than control. Taken in conjunction with our SHG imaging results, these observations allow us to build a pictorial model of collagen fibril pattern and its regulation by DS (Figure 7). Presence of lower levels of DS results in greater number of fibrils. However, a lower tendency to bundle and form higher order elements and lesser width and possibly, length results in poor mean SHG signals and surface occupancy. Presence of higher levels of DS restores ordering of fibrils, their width and length but the spatial density of fibrils formed is lower than in the control gels.

**Figure 6.**
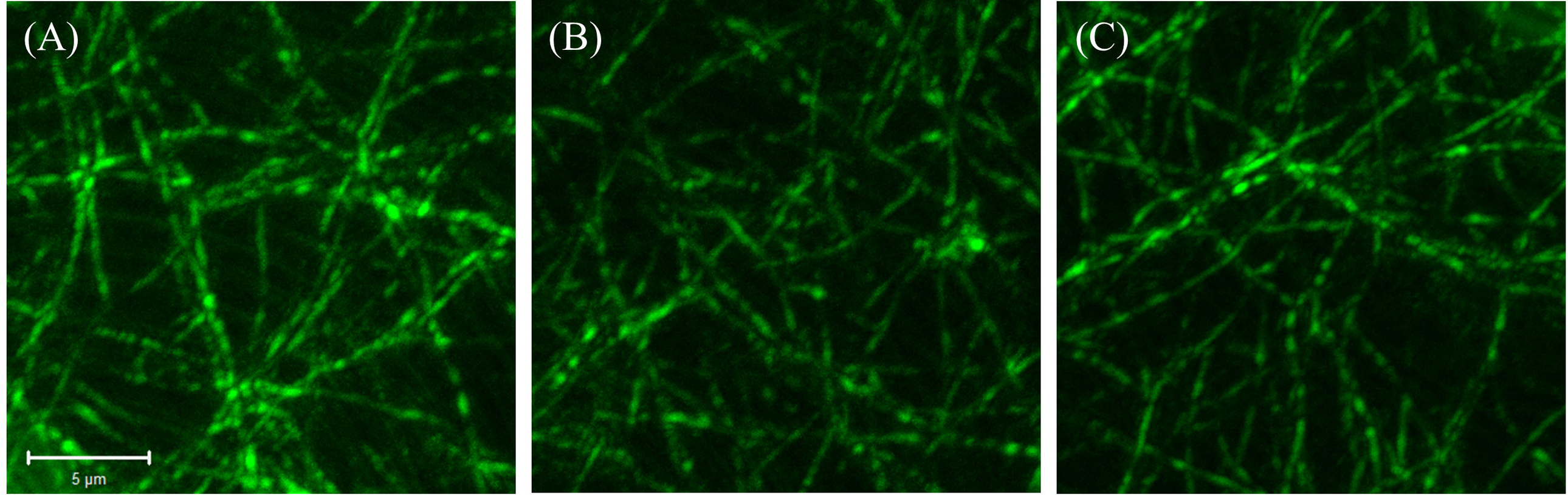
Confocal microscopic examination of Collagen I fibrils. Laser confocal reflectance micrographs of collagen I matrices polymerized in the absence of dermatan sulphate (A) and presence of dermatan sulphate at 50 µg/mL (B) and 400 µg/mL (C) respectively, showing reflectance signals emanating from the collagen fibrils post polymerization and fixation. Scale bar = 5 µm.

**Figure 7.**
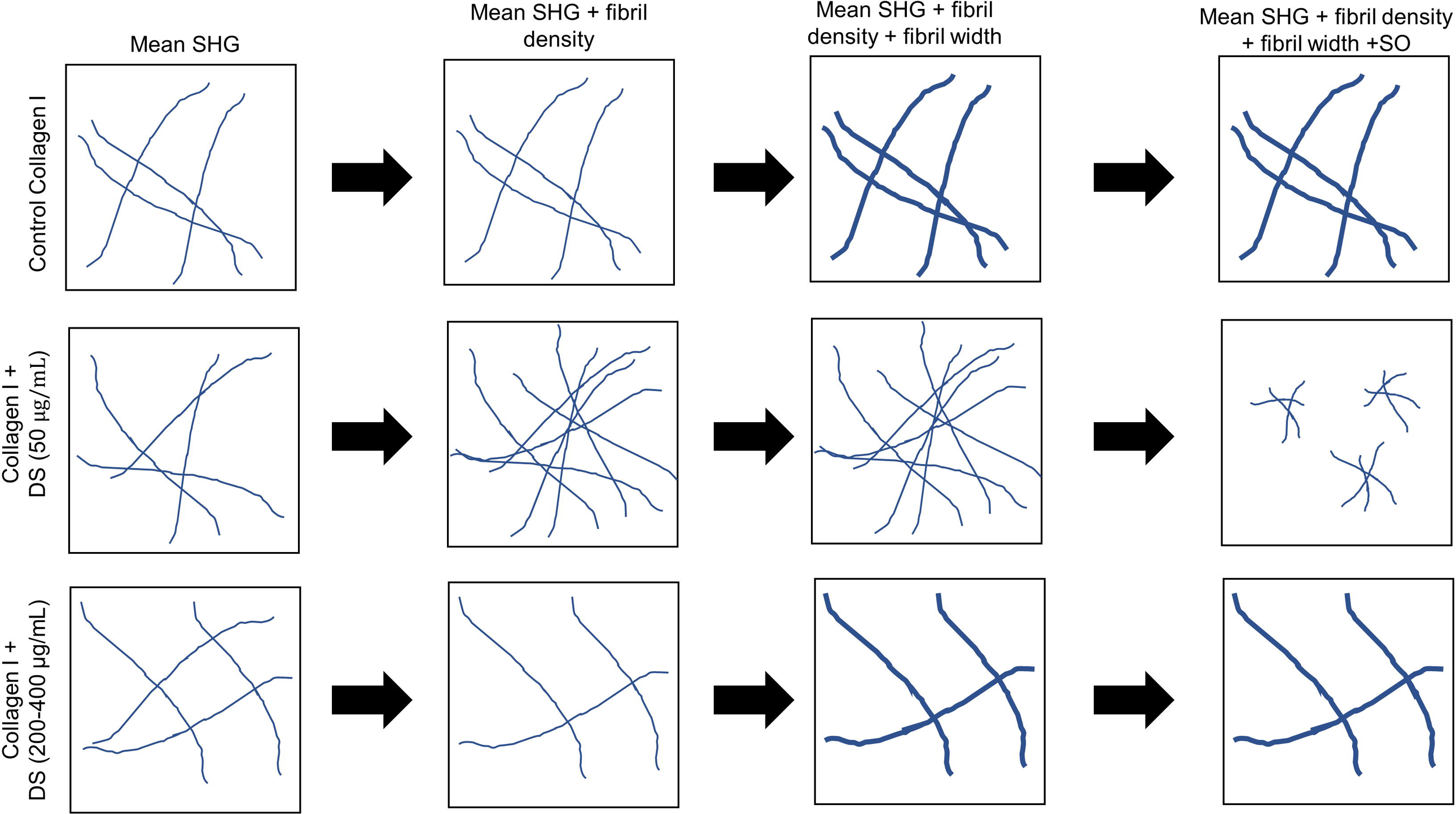
A schematic model of distinct collagen I gel architectures as an effect of differential levels of Dermatan Sulfate. Model depicting the reorganization of collagen fibrils in the absence (top row) and presence of DS at lower concentrations 50 µg/mL (middle row) and higher concentrations (200 µg/mL and 400 µg/mL (bottom row)). Columns represent the changes incorporated into the gel architecture with the gradational addition of each examined optical parameter.

## DISCUSSION

In 2018, an interdisciplinary team of clinicians and scientists sought to posit the interstitium of tissues as a potential new organ [40]. The proposition was based on the fact that histologically unprocessed organs and tissues consisted of interstitial spaces sculpted within the extracellular milieu containing ECM. Such interstitial spaces contribute to subtle functions that would otherwise not be detected when such tissues are fixed and dehydrated, resulting in loss of such spaces. The paper provided rich insights into the structure and function of interstitia using, among other microscopic techniques, second harmonic generation imaging. In fact, compressive forces acting on tissues during dehydration and sample processing may irrevocably alter the tissue microenvironment and distort the ECM architecture. This has necessitated the innovation of technologies that seek to image biological specimen in their native state as much as possible. In addition, second harmonic generation (SHG) has the advantage of being used to detect fibrillar structures such as collagen I in a mixture of heterogeneous biological materials.

Here we show that SHG imaging cannot just be used to detect Collagen I, but also is sensitive enough to detect subtle changes in its fibril arrangements that are wrought by the presence of different concentrations of Dermatan Sulfate (DS), a GAG that associates with it. In addition, our experiments help establish that the regulation of Collagen fibrillogenesis by DS to be a fairly complex process: the presence of high and low levels of DS during polymerization have distinct (fibril width, surface occupancy, ordering) and even opposite (fibril density) effects on Collagen I gel architecture. Why may this be the case? A careful reading of the literature on DS reveals that they have a propensity for self-aggregation [41, 42]. Therefore, the biophysical properties of high (and possibly aggregated) levels of DS are likely to be distinct from that of low (and dispersed) levels allowing them to impact Collagen I in unique ways. Our findings therefore indicate that lower levels of DS sculpt a Collagen I fibril pattern that is skewed towards lower ordered elements (less bunding and fibril formation). This explains lower mean SHG signals, lower element width and greater fibril number. In contrast, presence of higher DS levels biases the formation of higher ordered elements with greater width and higher mean SHG signals. The ratio of DS to Collagen levels we have used in this study correspond with tissue- and age-specific concentrations, although the changes of such ratios in pathological conditions remains to be elucidated [9, 43]. Our findings may therefore also shed light on how collagen fibril architectures can diverge so drastically between, for instance the cornea and the tendon, when the expression of DSPGs, especially decorin is thought to be ubiquitous [44, 45].

Our observations on DS-regulated Collagen I patterns are likely to be of value for synthesis of implant scaffolds, where different greater and weaker ordered architectures can now be tuned to customize a biocompatible scaffold that provides mechanical durability and/or acts as a depot to store diffusible molecules. The implications of our findings in the field of glycopathology are immense. DSPGs such as biglycans and decorin are known to be upregulated in several cancers. SHG imaging has been applied to investigate the morphological difference in normal and cancerous ex vivo human tissues [46]. A recent study also reports the use of SHG metrices such as emission directionality, relative conversion efficiency as well as wavelet transforms and texture analysis to demonstrate the effect of Decorin in organization of prostatic collagen matrix [47]. Image transforms of SHG micrographs such as curvelet transform and wavelet transforms have been used to identify tumor cell boundary required for Tumor-Associated Collagen Signature (TCS) to characterize the disease progression in breast cancer models [48]. Our study shows that the regulation of interstitial Collagen I architecture by such DSPGs may be much more complex than previously thought with higher and lower levels of DS influencing cancer cell motility and migration in divergent ways.

Future efforts will be directed at understanding the role of diverse DS levels on the manifestation and heterogeneity of cellular phenotype within collagenous microenvironments. Furthermore, SHG imaging can be combined with bond-specific vibrationally resonant sum-frequency generation spectroscopy [49] to understand the biochemical origins of modification of the asymmetric molecular units in collagen triple helix in the presence of proteoglycans. We would also study in detail how DS-modulated divergent Collagen I element architectures may exhibit distinct mechanical properties as has been elegantly shown albeit at a single DS concentration before [9]. This will further the potential for DS as a tuner of the architectural properties of the most abundant ECM protein of our body with wide implications for its future use in the field of bioengineering.

## Acknowledgments

RB and VR would like to acknowledge the Institute of Eminence grant (IE/CARE-19-0319) received from the Indian Institute of Science Bangalore for funding this research. In addition, RB would also like to acknowledge support from the Department of Biotechnology, India [BT/PR26526/GET/119/92/2017] and SERB [ECR/2015/000280]. VR would also like to acknowledge support from Department of Science & Technology through the Indo-Korea joint research projects (INT/Korea/P-44). PS would like to acknowledge funding support from DST INSPIRE fellowship (IF170197, IVR number-201600020331).

## References

1. Mouw, J.K., G. Ou, and V.M. Weaver, Extracellular matrix assembly: a multiscale deconstruction. Nat Rev Mol Cell Biol, 2014. 15(12): p. 771–85.

2. Theocharis, A.D., et al., Extracellular matrix structure. Adv Drug Deliv Rev, 2016. 97: p. 4–27.

3. Bonnans, C., J. Chou, and Z. Werb, Remodelling the extracellular matrix in development and disease. Nat Rev Mol Cell Biol, 2014. 15(12): p. 786-801.

4. Scott, J.E., Proteoglycan-fibrillar collagen interactions. Biochem J, 1988. 252(2): p. 313–23.

5. Yanagishita, M., Function of proteoglycans in the extracellular matrix. Acta Pathol Jpn, 1993. 43(6): p. 283–93.

6. Nash, M.A., M.T. Deavers, and R.S. Freedman, The expression of decorin in human ovarian tumors. Clin Cancer Res, 2002. 8(6): p. 1754–60.

7. Iozzo, R.V. and A.D. Murdoch, Proteoglycans of the extracellular environment: clues from the gene and protein side offer novel perspectives in molecular diversity and function. FASEB J, 1996. 10(5): p. 598–614.

8. Chen, S. and D.E. Birk, The regulatory roles of small leucine-rich proteoglycans in extracellular matrix assembly. FEBS J, 2013. 280(10): p. 2120–37.

9. Reese, S.P., C.J. Underwood, and J.A. Weiss, Effects of decorin proteoglycan on fibrillogenesis, ultrastructure, and mechanics of type I collagen gels. Matrix Biol, 2013. 32(7-8): p. 414–23.

10. Ameye, L. and M.F. Young, Mice deficient in small leucine-rich proteoglycans: novel in vivo models for osteoporosis, osteoarthritis, Ehlers-Danlos syndrome, muscular dystrophy, and corneal diseases. Glycobiology, 2002. 12(9): p. 107R–16R.

11. Kowitsch, A., G. Zhou, and T. Groth, Medical application of glycosaminoglycans: a review. J Tissue Eng Regen Med, 2018. 12(1): p. e23–e41.

12. Tierney, C.M., M.J. Jaasma, and F.J. O’Brien, Osteoblast activity on collagen-GAG scaffolds is affected by collagen and GAG concentrations. J Biomed Mater Res A, 2009. 91(1): p. 92–101.

13. Prestwich, G.D., Hyaluronic acid-based clinical biomaterials derived for cell and molecule delivery in regenerative medicine. J Control Release, 2011. 155(2): p. 193–9.

14. Vallen, M.J., et al., Primary ovarian carcinomas and abdominal metastasis contain 4,6-disulfated chondroitin sulfate rich regions, which provide adhesive properties to tumour cells. PLoS One, 2014. 9(11): p. e111806.

15. Li, H.P., et al., Roles of chondroitin sulfate and dermatan sulfate in the formation of a lesion scar and axonal regeneration after traumatic injury of the mouse brain. J Neurotrauma, 2013. 30(5): p. 413–25.

16. Junqueira, L.C. and G.S. Montes, Biology of collagen-proteoglycan interaction. Arch Histol Jpn, 1983. 46(5): p. 589–629.

17. Douglas, T., et al., Interactions of collagen types I and II with chondroitin sulfates A-C and their effect on osteoblast adhesion. Biomacromolecules, 2007. 8(4): p. 1085–92.

18. Gouignard, N., et al., Musculocontractural Ehlers-Danlos syndrome and neurocristopathies: dermatan sulfate is required for Xenopus neural crest cells to migrate and adhere to fibronectin. Dis Model Mech, 2016. 9(6): p. 607–20.

19. Mizumoto, S., et al., Pathophysiological Significance of Dermatan Sulfate Proteoglycans Revealed by Human Genetic Disorders. Pharmaceuticals (Basel), 2017. 10(2).

20. Nakamura, A., T. Osonoi, and Y. Terauchi, Relationship between urinary sodium excretion and pioglitazone-induced edema. J Diabetes Investig, 2010. 1(5): p. 208–11.

21. Appunni, S., et al., Small Leucine Rich Proteoglycans (decorin, biglycan and lumican) in cancer. Clin Chim Acta, 2019. 491: p. 1–7.

22. Bostrom, P., et al., Human Metaplastic Breast Carcinoma and Decorin. Cancer Microenviron, 2017. 10(1-3): p. 39–48.

23. Pavone, F.S. and P.J. Campagnola, Second harmonic generation imaging. 2016.

24. Masters, B.R. and P. So, Handbook of biomedical nonlinear optical microscopy. 2008.

25. Campagnola, P. and C.D. L, Laser and Photonics Review. 2011. 5(1).

26. IsAAcFREUND, MOSHEDEUTSCH, and ANDAARONSPRECHERt, Optical Second-harmonic Microscopy, Crossed-beam Summation, and Small-angle Scattering in Rat-tail Tendon. 1986.

27. Campagnola, P.J. and C.Y. Dong, Second harmonic generation microscopy: principles and applications to disease diagnosis. 2010.

28. Campagnola, P., Second harmonic generation imaging microscopy: applications to diseases diagnostics. Anal Chem, 2011. 83(9): p. 3224–31.

29. Rocha-Mendoza, I., et al., Sum Frequency Vibrational Spectroscopy: The Molecular Origins of the Optical Second-Order Nonlinearity of Collagen. 2007.

30. W, M., et al., Coherent nonlinear optical imaging: beyond fluorescence microscopy. 2011. 31.

31. Campbell, K.R. and P.J. Campagnola, Assessing local stromal alterations in human ovarian cancer subtypes via second harmonic generation microscopy and analysis. J Biomed Opt, 2017. 22(11): p. 1–7.

32. Watson, J.M., et al., Analysis of second-harmonic-generation microscopy in a mouse model of ovarian carcinoma. J Biomed Opt, 2012. 17(7): p. 076002.

33. Campbell, K.R., et al., Polarization-resolved second harmonic generation imaging of human ovarian cancer. 2018.

34. A, F., et al., Medical Image Computing and Computer-Assisted Interventation. 1998.

35. Xu, S., et al., Quantification of liver fibrosis via second harmonic imaging of the Glisson’s capsule from liver surface. 2016.

36. Otsu, N., A Tlreshold Selection Method from Gray-Level Histograms. 1979.

37. Glaser, M., et al., Self-assembly of hierarchically ordered structures in DNA nanotube systems. New Journal of Physics, 2016. 18(5).

38. C, S., An unbiased detector of curvilinear structures. 1998.

39. Xiyi Chen, et al., Second harmonic generation microscopy for quantitative analysis of collagen fibrillar structure.

40. Benias, P.C., et al., Structure and Distribution of an Unrecognized Interstitium in Human Tissues. Sci Rep, 2018. 8(1): p. 4947.

41. Scott, P.G., et al., Proteoglycans of the articular disc of the bovine temporomandibular joint. II. Low molecular weight dermatan sulphate proteoglycan. Matrix, 1989. 9(4): p. 284–92.

42. Coster, L., et al., Self-association of dermatan sulphate proteoglycans from bovine sclera. Biochem J, 1981. 197(2): p. 483–90.

43. McKee, T.J., et al., Extracellular matrix composition of connective tissues: a systematic review and meta-analysis. Sci Rep, 2019. 9(1): p. 10542.

44. Mauviel, A., et al., Transcriptional regulation of decorin gene expression. Induction by quiescence and repression by tumor necrosis factor-alpha. J Biol Chem, 1995. 270(19): p. 11692–700.

45. Trowbridge, J.M. and R.L. Gallo, Dermatan sulfate: new functions from an old glycosaminoglycan. Glycobiology, 2002. 12(9): p. 117R–25R.

46. Chen, X., et al., Quantitative analysis of collagen change between normal and cancerous thyroid tissues based on SHG method, in Tenth International Conference on Photonics and Imaging in Biology and Medicine (PIBM 2011). 2012.

47. Campbell, K.R., et al., Second-harmonic generation microscopy analysis reveals proteoglycan decorin is necessary for proper collagen organization in prostate. J Biomed Opt, 2019. 24(6): p. 1–8.

48. Tilbury, K. and P.J. Campagnola, Applications of second-harmonic generation imaging microscopy in ovarian and breast cancer. Perspect Medicin Chem, 2015. 7: p. 21–32.

49. Raghunathan, V., et al., Rapid vibrational imaging with sum frequency generation microscopy. Opt Lett, 2011. 36(19): p. 3891–3.

