## Supplementary figures and images for "Differential levels of dermatan sulfate generate distinct Collagen I gel architectures"

### Supplementary Figure1

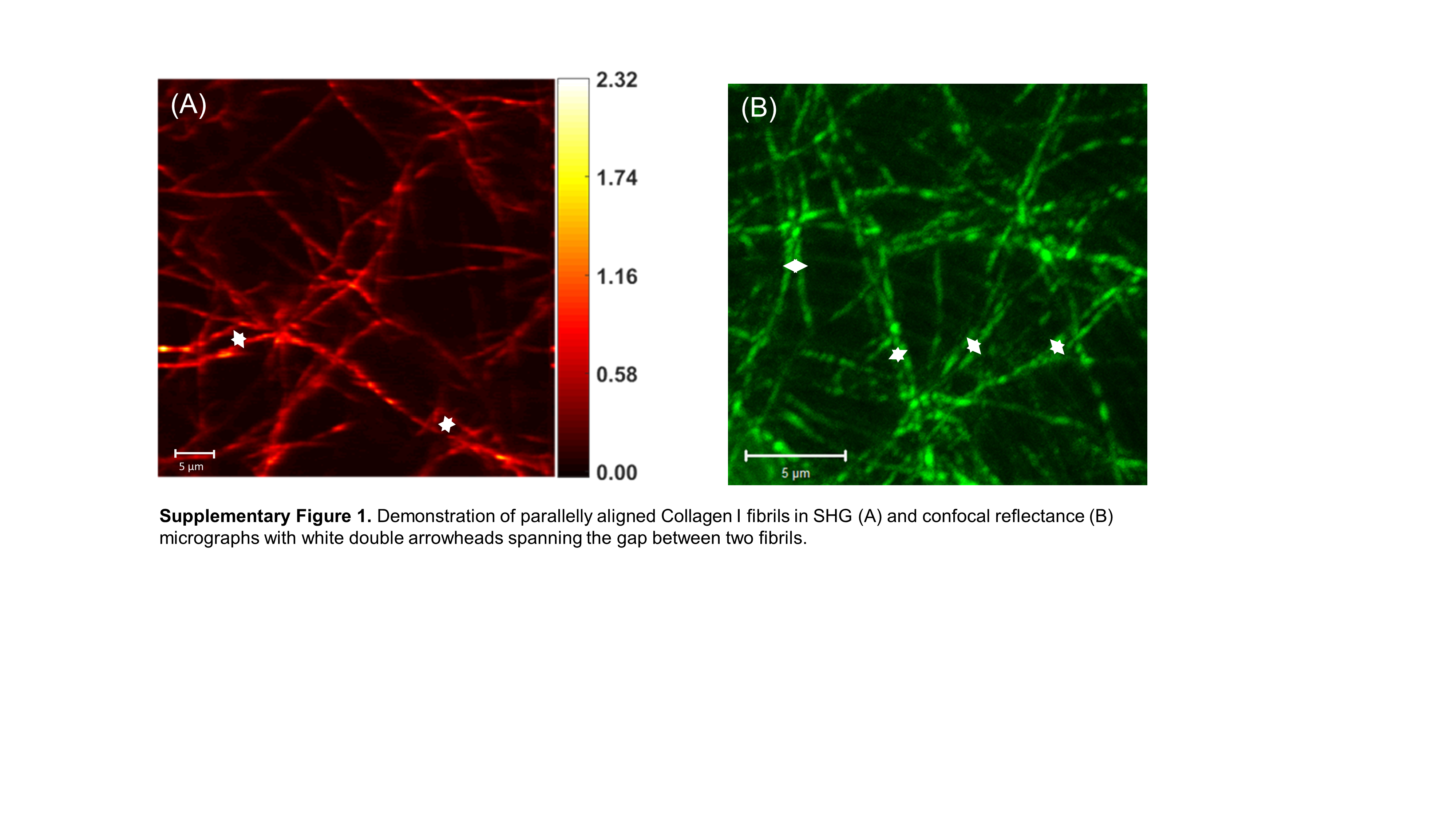

### Supplementary Figure 2

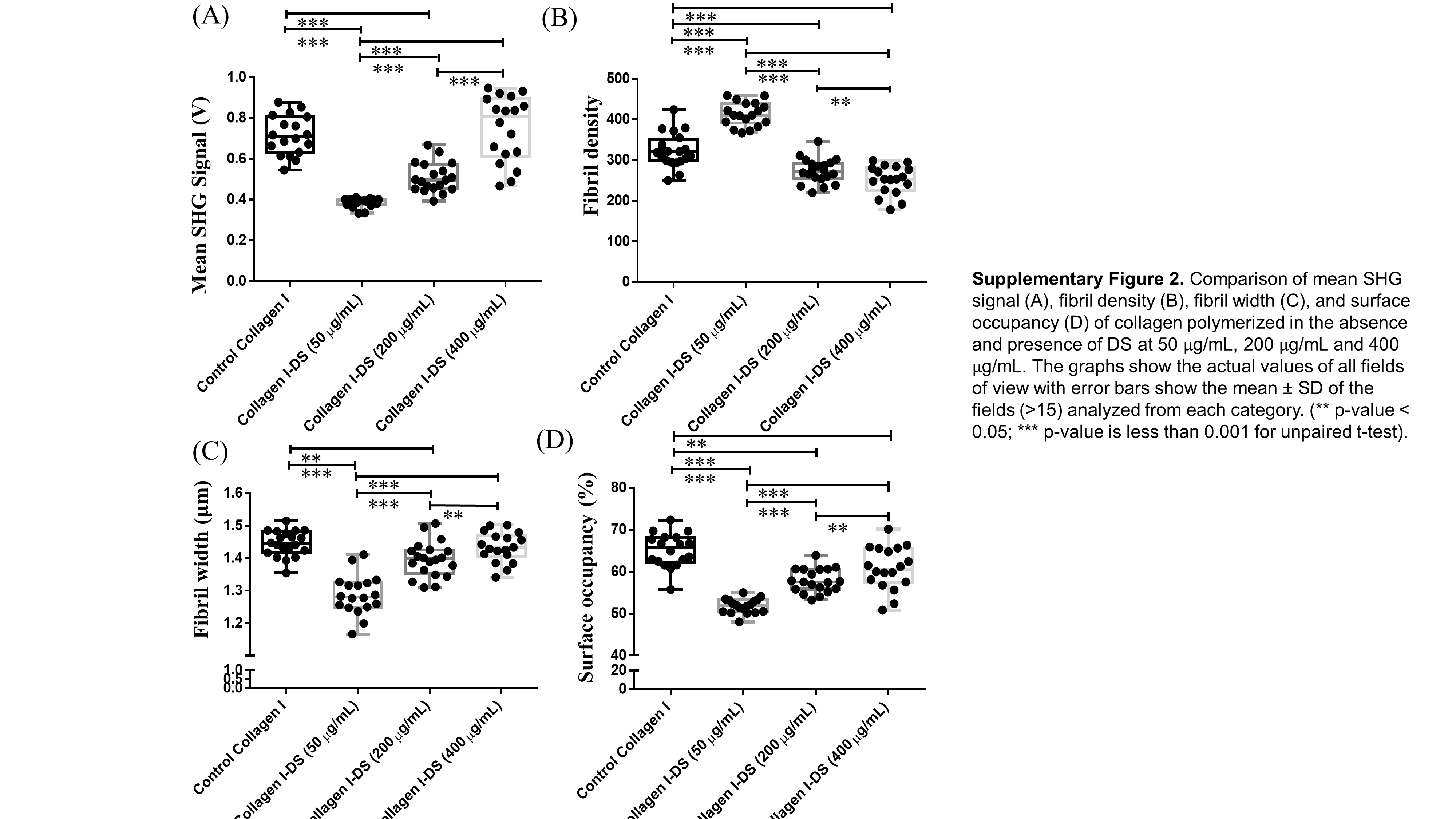
